# IgG derived dendritic cells can induce production of IL-17 by T cells in multiple sclerosis

**DOI:** 10.1101/2022.08.15.503963

**Authors:** Nazanin Pournasrolla, Ehsan Ahmadi, Seyedbahaadin Siroos, Maryam Nourizadeh, Mohammad Hossein Harirchian, Maryam Izad

## Abstract

Multiple sclerosis (MS) is an autoimmune demyelinating disease of the central nervous system (CNS). Myelin-autoreactive T cells have been implicated in the initiation of an inflammatory cascade. Dendritic cells (DC) are key modulators of this immuno-pathological cascade. The interaction between immune complexes (IC) and FcγRs results in activation of the immune system and induction of host inflammatory responses. Otherwise, monocytes differentiate into DCs after ligation of their FcγRs to IgG. We investigated circulating immune complexes levels (CIC) and differentiation of monocytes onto immature dendritic cell (iDC) via FcγR by Plate-bound human IgG in MS patients compared to healthy individuals. Our results showed that the concentration of CIC in patients with MS was significantly higher than healthy controls. Human IgG alone differentiate monocytes into DCs with a phenotype, including up-regulation of CD1b, CD86 and down-regulation of CD14. Also, the ability of LPS/MBP matured DCs in activation and cytokine production of autologous T cells was evaluated by MLR assay and ELISA. The level of IL-17 was significantly higher in MS patients when IgG derived DCs cocultured with T cells. Also, a correlation between IL-17 levels and circulating immune complexes level was observed in MS patients. Therefore, activation of FcγR on monocytes triggers differentiation into specialized iDC with the capacity to induce auto-reactive T cells that may contribute to the pathogenesis of MS.

## Introduction

Multiple sclerosis (MS) is an autoimmune disease that causes demyelination of the nerves of central nervous system (CNS) (1). MS is a complex disease and many risk factors such as vitamin D deficiency, ultraviolet (UV) light exposure, Epstein-Barr virus (EBV) infection, smoking, obesity and genetic factors (like HLA genes) increase disease susceptibility (2).

MS is considered to be a T cell-mediated autoimmunity. Pro-inflammatory Th1 and Th17 cells involve in the pathogenesis of MS and Experimental Autoimmune Encephalomyelitis (EAE), which is a widely used animal model of MS (3, 4).The activity of these T cells is modulated by other cell populations, specifically with dendritic cells. Dendritic cells, like B cells and macrophages, are professional antigen presenting cells (APCs). However, DCs are more powerful than other APCs in inducing the differentiation of naïve CD4^+^ T cells into subclasses of T helper cells. Also, dendritic cells have unique abilities in antigen processing, expression of co-stimulatory molecules, cytokine production, and migration to inflamed organs or lymphoid tissues to prime T cells (5, 6). Therefore, DCs are critical for initiation and development of T cell inflammatory responses in inflamed organs such as CNS of MS patients.

Dendritic cells as a heterogeneous population are divided into four classes: classical dendritic cell-1 (cDC1), cDC2, plasmacytoid dendritic cell (pDC), and monocyte-derived dendritic cells (mo-DC) based on differentiation origin, surface markers, and function (7). cDC1 and cDC2 mostly activate Th1 and Th2 responses, respectively. On the other hand, pDCs play a critical role in induction of anti-viral immune responses by producing large amount of type 1 interferons (8). The mo-DCs differentiate from monocyte and express CD11, MHC II, FCγRI, CD14, FCεRI, CD1a/CD1c and CD172a (SIRPα) markers (9-11). Mo-DCs also known as inflammatory DCs because they accumulate in inflamed tissues. Although the exact role of mo-DCs has not defined yet, it is widely accepted that mo-DCs have a synergic with cDCs in response to inflammation or infection (9, 12). Therefore, further studies are needed to investigate the role of mo-DCs during inflammatory conditions namely in autoimmune responses.

FCγ receptors (FCγR) broadly express on the surface of immune cells such as macrophages, DCs, neutrophils (13). In human, there are three types of FCγ receptors: FCγRI, FCγRIIA/B, and FCγRIIIA/B. Between them, FCγRI, FCγRIIA, and FCγRIII have activating role and FCγRIIB has inhibitory role (14). Human and mouse dendritic cells, express a wide spectrum of FCγ receptors, which can describe the potential role of these cells in antigen presentation through interaction with IgG immune complexes (15). Furthermore, activation of monocytes via FCγR can trigger differentiation into DCs (16-18). Also, increased expression of FcγRs on the surface of microglial cells have been shown in active lesion of MS, which mediated phagocytosis, antibody-dependent cytotoxicity and released oxygen radicals (19). Otherwise, activation of FcγRs on the surface of monocytes can trigger differentiation into specialized iDCs with excessive potential to induce auto-reactive T cells that may contribute to the pathogenesis of MS. Also, there are other methods for differentiation of monocytes into mo-DCs. Culture of monocytes with both GM-CSF and IL-4 providing a model for the in vitro generation of DCs from monocytes to investigate the effect of various agents on the differentiation pathways (20, 21). Triggering differentiation of monocyte into mo-DCs with immune complexes is another helpful way to study mo-DCs function because different immune complexes have been observed in the sample of patients with autoimmunity that have potential for induction of mo-DCs (22, 23).

An increasing number of studies search for the role of DCs in induction and maintenance of autoimmunity, such as MS. However, there is no study examining the role of IgG mediated immune complexes in differentiation of iDCs to induce auto-reactive T cells in MS patients. Hence, we decided to evaluate the differentiation of monocytes into mo-DC using immobilized human IgG and also function of T cells after recognition of MBP on iDCs in MS and healthy samples.

## Materials and methods

### MS patients and Healthy volunteers

Twenty patients (3 male,17 females, aged 22-45 years, median age 33) with definite relapsing– remitting multiple sclerosis (RRMS) according to McDonald’s criteria (24) and 20 healthy volunteers (3 male,17 females, aged 24 - 44 years, median age 34) were enrolled in this study. None of the patients were under immunosuppressive or immune-modulatory therapies at least three months before the sampling. Healthy controls had no past family history of MS or other autoimmune diseases. This research was approved by ethics committee of the Tehran University of Medical Sciences and written informed consent was obtained from all participants. All of the patients were referred to Iranian Center of Neurological Research in Imam Khomeini General Hospital, Tehran University of Medical Sciences, Tehran, Iran.

### Differentiation of monocyte to DCs

Fresh Peripherial Blood Mononuclear Cells (PBMCs) were isolated from heparinized venous blood by density gradient centrifugation over Ficoll Lymphodex (inno-Train, Diagnostic Gumby, Germany). Highly enriched blood monocytes (90-95% CD14+) were obtained using anti-CD14 Micro Beads kit (Miltenyi Biotic cat. No. 130-050-21, Germany) according to the kit instructions. To obtain IgG derived DCs, culture plates were first coated by various concentrations of human (h) IgG derived from healthy donor-pooled sera (25, 100, 500µg/ml into PBS) and washed twice with PBS. before monocyte cultures.

Monocytes were cultured at 37°C in RPMI 1640 supplemented with penicillin (100 IU/ml) and streptomycin (100 µg/ml), 1% L-glutamine (100 mM), 1% sodium pyruvate (1M) (Sigma, US) and 10% fetal calf serum (Gibco, Thermo Fisher Scientific, US). The monocytes were stimulated with medium, immobilized hIgG, or 50 ng/ml recombinant GM-CSF (rGM-CSF) (Bioscience, US) plus 20 ng/ml IL-4 (R&D systems Cat.No. 204-IL-010, US), for 5 days to obtain immature dendritic cells (iDC). Maturation of iDCs was performed in the presence of 200 ng/ml LPS from *Escherichia coli* (Sigma Aldrich, Germany) and 10µg/ml myelin basic protein (MBP) (87-99 residues) (ANA SPEC Cat.No.29211, US) for 48 hours.

### Flow cytometric analysis

For phenotypic analysis of immature and mature DCs, cells were harvested on days 5 and 7. The cells were stained by anti-Human CD1b-APC, CD116-FITC, CD86-PE, CD14-PerCP-Cy5.5, CD83-FITC, CD40PE-Cy™5 and appropriate isotype controls (BD Biosciences, US) for 30 min at 4°C. Cells were analyzed with a FACSCalibur flow cytometer (BD Bioscience) (BD Biosciences, US) using Cell Quest software. Finally, exported data were analyzed with FlowJo™ v10.5.3 software (BD Life Sciences).

### Evaluation of T cell responses

We isolated T cells from PBMC of patients and healthy controls for autologous mixed lymphocyte reaction (MLR) assay using Dynabeads Untouched Human CD4^+^ T Cells kit (Invitrogen Dynal cat. No.113.52D, US). For MLR, FcγR-activated and rGM-CSF+IL-4 induced DCs (2×10^4^ cells/ml) were stimulated by LPS or MBP, and co-cultured with 2×10^5^ T cells for 72 hours.

### Measurement of cytokines and Immune Complexes

Supernatants from DC cultures on days 5, 7, and after co-culturing with T cells were collected and stored at -70°C until testing for the presence of cytokines. We measured IFN-γ, IL-17, IL-12p70 (R&D Systems, US) and IL-23 (BD Biosciences, US) levels in supernatants using ELISA (BD Biosciences and R&D Systems, US) according to kits instructions. We also measured CIC levels in serum of patients and controls by Anti-C1q CIC assay (Dia Metra, DKO016, Italy).

### Statistical analysis

Data are expressed as mean ± SD. Statistical analyses for parametric data were performed with One-way ANOVA test and for non-parametric data, K Independent Samples test were used. P values less than 0.05 were considered statistically significant between groups. All analysis were performed using the SPSS v.22 software (IBM).

## Results

### Circulating immune complexes (CIC) increased in MS patients

A total number of 40 subjects (20 RRMS patients and 20 healthy controls) were enrolled in this study. Sera of patients and controls were evaluated for circulating immune complexes levels by Anti-C1q ELISA method. Our results showed that the concentration of CIC in patients with MS was significantly higher than healthy controls (P value < 0.001) (Fig.1, a).

**Figure 1.**
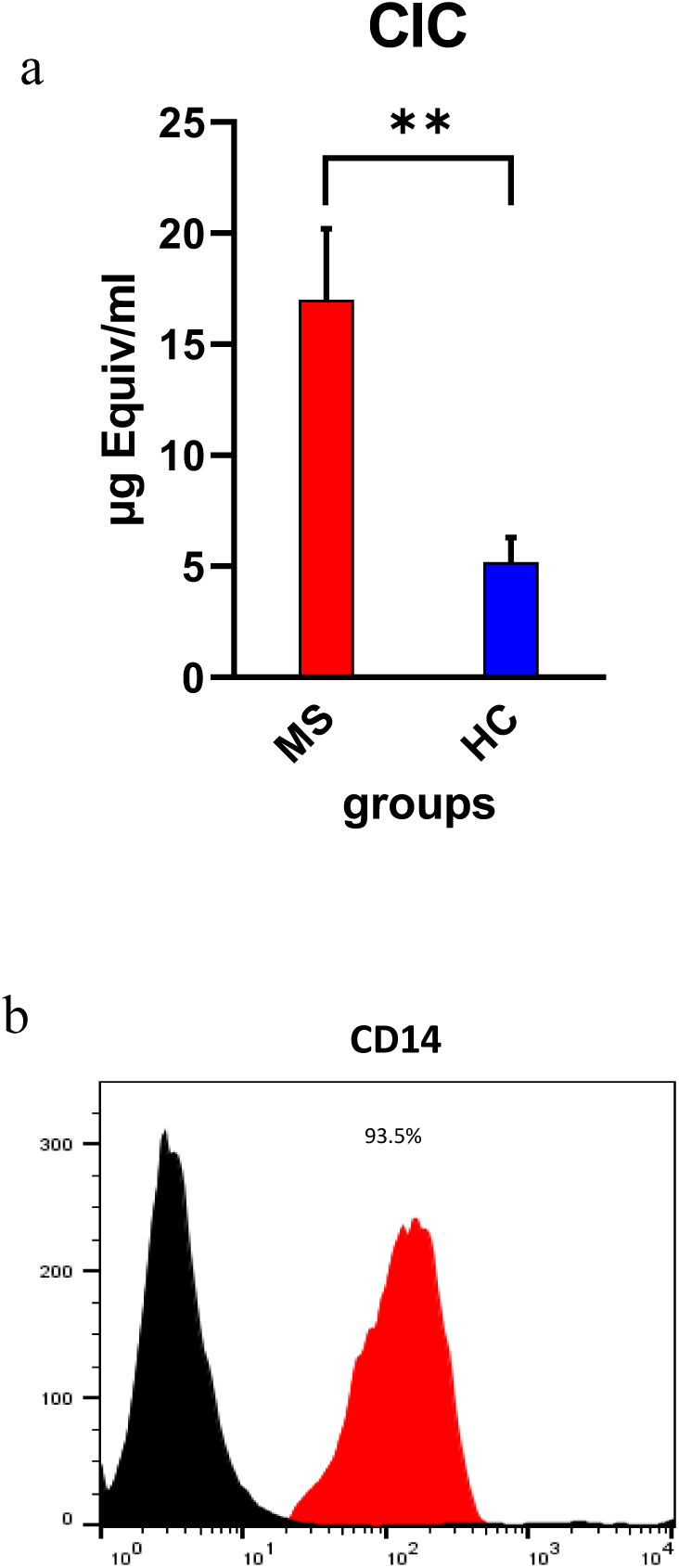
The results of evaluating CIC in sera of MS patients and healthy controls by Anti-C1q method are shown as bar graph. The result of purification of monocytes showed by histogram as the percentage of CD14+ cells (b). The significancy of results are shown as *<0.05, **<0.01, and ***<0.001.

### FcγR stimulation with IgG caused monocyte differentiation into immature DCs

In this study we found that the concentration of CIC increased in MS patients. Otherwise, Tanaka et al. (25) showed that immune complexes can differentiate monocytes into dendritic cells and these cells can induce auto-reactive T cells. Regarding to these results, we examined the phenotypic and functional differences of IgG or GM-CSF monocyte-derived dendritic cells between MS patients and controls.

Highly enriched blood monocytes (90-95% CD14^+^) (Fig.1, b) isolated from the peripheral blood were cultured in various concentrations (25, 100 and 500 µg/ml) of immobilized human serum IgG (Fig.2, a). The result showed that immobilized IgG in concentration of 100 µg/ml has greater potential for differentiation of monocytes to DCs (CD1b^+^) compared to other concentrations. Therefore, we used this concentration for next experiments of our study.

**Figure 2.**
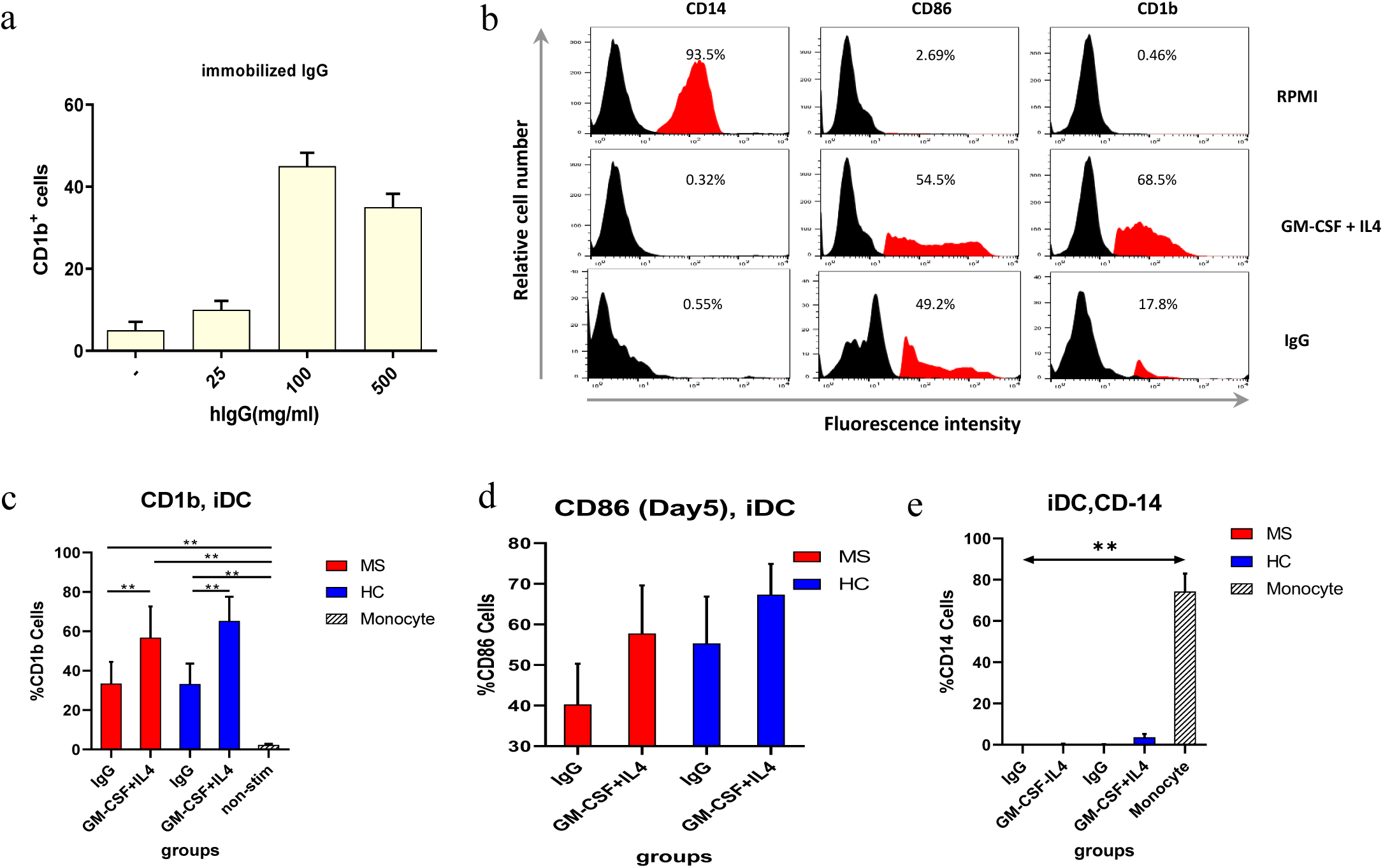
Monocytes were stimulated with various concentrations of immobilized IgG (hIgG), five days later, the percentage of CD1b^+^ cells was determined by flow cytometry (a). Human monocytes were stimulated with hIgG (100 µg/ml), or rGM-CSF (50ng/ml)+ IL-4 (20 ng/ml), or medium (un-stimulated). After 5 days, expression of cell surface markers were examined by flow cytometry. Representative histograms for CD14, CD86, and CD1b are shown to illustrate the differentiation stage of the DC (b). The statistical comparison of expression of these markers across different groups is shown by bar graphs by mean + SEM (c, d, e). The significancy of results are shown as * <0.05, **<0.01, and ***<0.001.

Monocytes were cultured in the presence of rGM-CSF plus IL4 combination or IgG alone. On day 5, cells were harvested and the expression of CD1b, CD14, and CD86 molecules on the surface of FcγR or GM-CSF+IL-4 derived-iDCs were analyzed by flow cytometry. The expression patterns of cell surface markers for both FcγR and rGM-CSF+IL-4 treated monocytes were consistent with an iDC-like phenotype, including high levels of CD1b, CD86 (B7.2, a co-stimulatory molecule for activation of T cells), as well as low expression of CD14 (Fig. 2, b).

The level of CD1b expression was found to be significantly higher on rGM-CSF+IL-4 iDCs compared to IgG derived iDCs (Fig. 2, c). Although the CD86 expression was higher in GM-CSF+IL-4 derived iDCs compared to IgG derived iDCs, it was not significant between groups (Fig. 2, d and e). Taken together, these data suggest that bound hIgG similar to rGM-CSF+IL-4 induces monocyte differentiation to iDCs.

### GM-CSF receptor mediates the induction of iDCs by FcγR from monocytes

Tanaka et al (25) showed that CD16-Cd64+ monocytes express higher levels of CD116 (α-chain of GM-CSF receptor) and hIgG activates these cells via CD64, results in upregulation of GM-CSF and differentiation of monocytes into CD1b iDCs. In this regard, Our question was whether the expression of CD 116 on iDC of MS patients was higher than healthy individuals. Therefore, we evaluated expression levels of CD116 on these cells (Fig. 3, a). We found the same expression level of GM-CSF receptors on FcγR derived iDCs versus rGM-CSF+IL-4 iDCs. There was no difference between patients and controls (Fig. 3, b).

**Figure 3.**
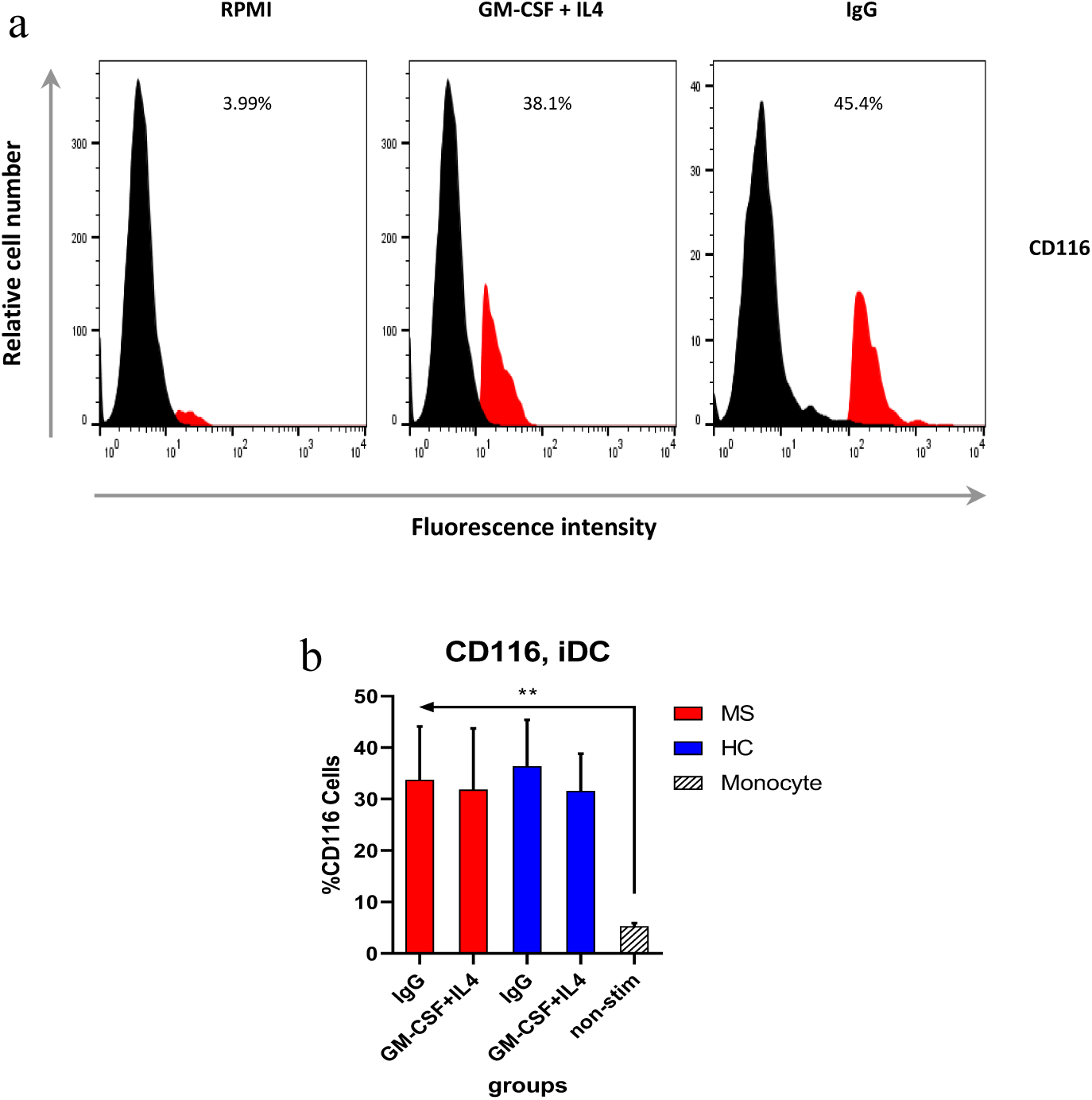
Human monocytes were stimulated with hIgG (100µg/ml) or rGM-CSF (50ng/ml) + IL-4 (20 ng/ml). After 5 days, expression of cell surface marker CD116 was examined by flow cytometry (a). Data represents mean percentage of CD116 expression + SEM (b). The significancy of results are shown as *<0.05, **<0.01, and ***<0.001.

### Maturation of iDCs by LPS and MBP and evaluation of T cell responses

Both FcγR and rGM-CSF+IL-4 derived iDCs were stimulated with LPS and MBP. After 48 hours incubation, the expression of costimulatory molecules, such as CD83, CD86, and CD40, as DC maturation markers were measured (Fig 4, a). The expression of both CD83 and CD40 markers were significantly higher in rGM-CSF+IL-4 derived DCs compared to IgG derived DCs. Also these markers were higher in MS samples compared to healthy samples (Fig. 4, b and c). Similar to iDC results, the CD86 expression in mature DCs was higher in GM-CSF+IL-4 treated DC than IgG induced DCs in both healthy and MS groups. CD86 expression in mature DCs showed no statistically significant result (Fig. 4d).

**Figure 4.**
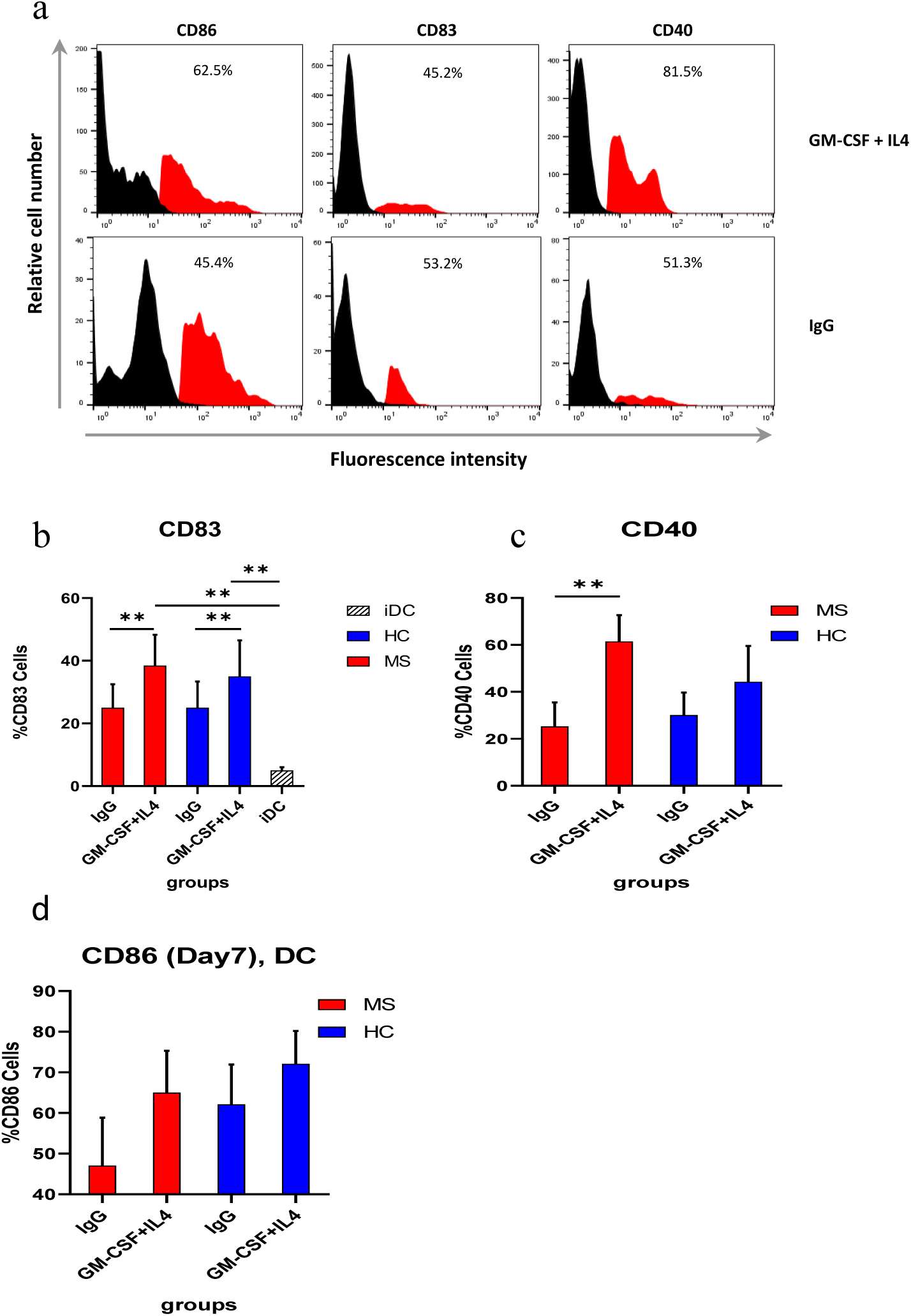
Both iDC groups were stimulated with LPS and MBP. After 48 hours, expression of cell surface markers CD83, CD86, and CD40 were examined by flow cytometry. Representative histograms are shown to illustrate the differentiation stage of the DC (a). The statistical comparison of expression of these markers across different groups is shown by bar graphs by mean + SEM (b, c, d). The significancy of results are shown as *<0.05, **<0.01, and ***<0.001.

Finally, the ability of both DCs in activation and cytokine production of autologous T cells was evaluated by MLR assay and ELISA. The levels of IL-17 were significantly higher in IgG derived DC compared to GM-CSF+IL-4 derived DCs in both MS samples and healthy controls (Fig. 6, a). Also, the correlation between IL-17 levels and circulating immune complexes level in MS patients were observed (Spearman’s rho = 0.053) (P = 0.04) (Fig. 6, b).

The result of evaluation of IFN-γ production of T lymphocyte in confronting to mature DCs showed significant lower level in MS patients compared to healthy controls. Moreover, the IFN-γ levels were identical between IgG and GM-CSF+IL-4 groups (Fig. 6, c). No correlation between IFN-γ production and circulating immune complexes level in RRMS patients were observed (Fig. 6, d).

On the other hand, the both DC groups were also investigated for cytokine production profile after maturation with LPS and MBP. The assessment of IL-23 production of DCs showed no significant differences between groups of study (Fig. 5, a). GM-CSF+IL-4 iDCs of MS patients significantly produced more IL-12 than those in healthy controls (P values < 0.05). IgG derived mature DCs also produced high levels of IL-12 although it was not significant (Fig. 5, b).

**Figure 5.**
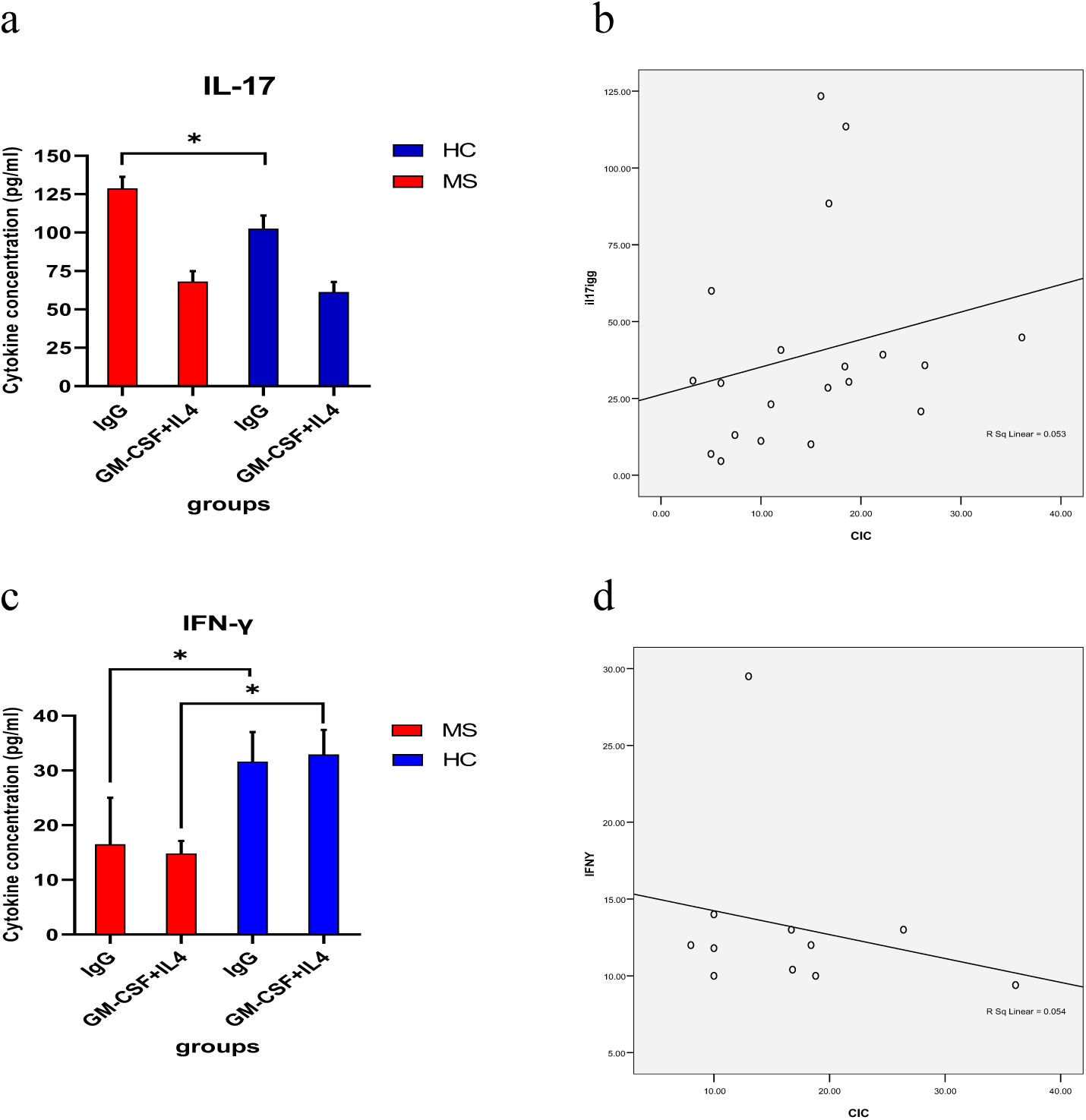
The ability of both DC groups in activation and cytokine production of autologous T cells was evaluated by MLR assay and ELISA. Supernatants were tested for presence of IL-17 (a, b) and IFN-γ (c, d). The results are shown as bar graph by mean + SEM. The significancy of results are shown as *<0.05, **<0.01, and ***<0.001.

**Figure 6.**
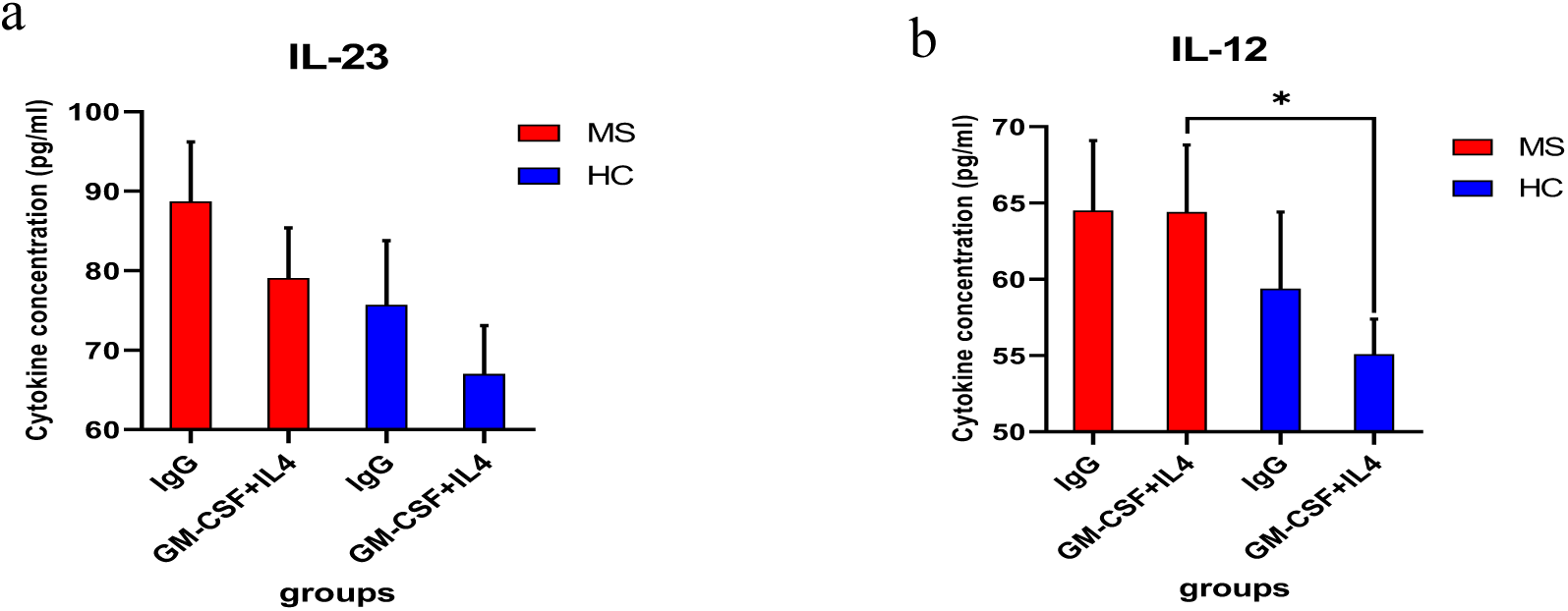
Supernatant from DC cultures were tested for presence of IL-12p70 and IL-23 cytokines by ELISA (a, b). The results are shown as mean + SEM. The significancy of results are shown as *<0.05, **<0.01, and ***<0.001.

## Discussion

Multiple sclerosis is considered to be a predominantly T cell-mediated disease. Evidences indicate that dendritic cells also have a critical role in the initiation and progression of the disease (26, 27). During inflammation and infection, dendritic cells derived from monocyte precursors by cytokines, including granulocyte-macrophage colony stimulating factor (GM-CSF) and macrophage colony stimulating factor (M-CSF), are mobilized towards inflammation sites (28, 29). MS patients seem to have higher levels of myeloid dendritic cells in their blood. Furthermore, dendritic cells derived from blood monocytes of MS patients secrete higher levels of pro-inflammatory cytokines than the same cells taken from healthy individuals (19, 30, 31). Previously, signaling through FcγR is known to inhibit the differentiation and maturation of cytokine derived DCs (32). Studies demonstrated that FcγR activation of human peripheral blood monocytes induces differentiation into CD1^+^ immature DCs with the potential to promote autoreactive T cell responses (25, 33).

In this study, we also showed that FcγR activation of peripheral blood monocytes induces differentiation into iDCs and these iDCs can stimulate autologous myelin specific T cells in MS patients similar to healthy controls. We found that the levels of IL-17 in the supernatant of IgG derived DC and T cells cocultures were significantly higher than GM-CSF+IL-4 derived DCs in both MS samples and healthy controls. In addition, the level of IL-17 was significantly higher in MS patients compared to controls.

Phenotypic and functional deficiencies of mono-DC have been reported in patients affected with autoimmune diseases (34). Mo-DCs from systemic lupus erythematosus (SLE) patients cultivated on immobilized IgG have shown considerably impaired maturation capability. These findings suggest that immune complexes might involve in the regulation of DC maturation (35). The present study provides evidences that FcγR cross-linking triggers differentiation of monocytes into iDCs and these iDCs stimulate autologous T cells in MS patients.

Although the combination of GM-CSF+IL-4 result in the generation of potent monocyte derived DCs that are more efficient, we have shown that FcγR activated DCs were comparable with these DCs in terms of markers and functions.

Immune complex formation is a response of immune system in situation of inflammation. The concentration of CIC in sera of rheumatoid arthritis (RA), SLE, or systemic scleroderma (SSc) patients was significantly higher comparing to healthy persons’ sera (36). Previous study demonstrated that increased level of auto-antibodies and immune complexes associated with exacerbation of MS in patients (19, 22). Consistent with other studies, our results showed that the immune complex level increased in RRMS patients in acute phase. Interestingly, there was a correlation between circulating immune complexes and IL-17 levels in MS patients. These ICs may have potentiality of differentiation of monocytes into iDCs via FcγRs.

Expression of FcγRs on the surface of microglia and macrophage cells has been increased in MS. Via these FcγRs, microglia and macrophage can mediate important effector functions that finally causes immune mediated demyelination and axonal degeneration during MS (37). Our findings in this study reveal a new level of immune response in immune-pathogenesis of MS through the interaction of IC with FcγR during the differentiation and maturation of dendritic cells. These findings may help to better elucidate the emerging role of DCs in MS pathogenesis and also to develop novel immunotherapeutic strategies for the treatment of this autoimmune disease.

## References

1. Dobson R, Giovannoni G. Multiple sclerosis - a review. Eur J Neurol. 2019;26(1):27–40.

2. Olsson T, Barcellos LF, Alfredsson L. Interactions between genetic, lifestyle and environmental risk factors for multiple sclerosis. Nature Reviews Neurology. 2017;13(1):25–36.

3. Etesam Z, Nemati M, Ebrahimizadeh MA, Ebrahimi HA, Hajghani H, Khalili T, et al. Altered Expression of Specific Transcription Factors of Th17 (RORγt, RORα) and Treg Lymphocytes (FOXP3) by Peripheral Blood Mononuclear Cells from Patients with Multiple Sclerosis. J Mol Neurosci. 2016;60(1):94–101.

4. Raphael I, Nalawade S, Eagar TN, Forsthuber TG. T cell subsets and their signature cytokines in autoimmune and inflammatory diseases. Cytokine. 2015;74(1):5–17.

5. Comabella M, Montalban X, Münz C, Lünemann JD. Targeting dendritic cells to treat multiple sclerosis. Nature Reviews Neurology. 2010;6(9):499–507.

6. Steinman RM. Dendritic cells: versatile controllers of the immune system. Nature Medicine. 2007;13(10):1155–9.

7. Guilliams M, Ginhoux F, Jakubzick C, Naik SH, Onai N, Schraml BU, et al. Dendritic cells, monocytes and macrophages: a unified nomenclature based on ontogeny. Nature Reviews Immunology. 2014;14(8):571–8.

8. Imai J, Otani M, Sakai T. Distinct Subcellular Compartments of Dendritic Cells Used for Cross-Presentation. Int J Mol Sci. 2019;20(22).

9. Collin M, Bigley V. Human dendritic cell subsets: an update. Immunology. 2018;154(1):3–20.

10. Collin M, Bigley V, Haniffa M, Hambleton S. Human dendritic cell deficiency: the missing ID? Nat Rev Immunol. 2011;11(9):575–83.

11. Eisenbarth SC. Dendritic cell subsets in T cell programming: location dictates function. Nature Reviews Immunology. 2019;19(2):89–103.

12. Auffray C, Fogg DK, Narni-Mancinelli E, Senechal B, Trouillet C, Saederup N, et al. CX3CR1+ CD115+ CD135+ common macrophage/DC precursors and the role of CX3CR1 in their response to inflammation. J Exp Med. 2009;206(3):595–606.

13. Boruchov AM, Heller G, Veri MC, Bonvini E, Ravetch JV, Young JW. Activating and inhibitory IgG Fc receptors on human DCs mediate opposing functions. J Clin Invest. 2005;115(10):2914–23.

14. Nimmerjahn F, Ravetch JV. Fcγ receptors as regulators of immune responses. Nature Reviews Immunology. 2008;8(1):34–47.

15. Bruhns P, Iannascoli B, England P, Mancardi DA, Fernandez N, Jorieux S, et al. Specificity and affinity of human Fcgamma receptors and their polymorphic variants for human IgG subclasses. Blood. 2009;113(16):3716–25.

16. Bánki Z, Kacani L, Müllauer B, Wilflingseder D, Obermoser G, Niederegger H, et al. Cross-linking of CD32 induces maturation of human monocyte-derived dendritic cells via NF-kappa B signaling pathway. J Immunol. 2003;170(8):3963–70.

17. Krutmann J, Kirnbauer R, Köck A, Schwarz T, Schöpf E, May LT, et al. Cross-linking Fc receptors on monocytes triggers IL-6 production. Role in anti-CD3-induced T cell activation. J Immunol. 1990;145(5):1337–42.

18. Krutzik SR, Tan B, Li H, Ochoa MT, Liu PT, Sharfstein SE, et al. TLR activation triggers the rapid differentiation of monocytes into macrophages and dendritic cells. Nat Med. 2005;11(6):653–60.

19. Vedeler CA, Myhr KM, Nyland H. Fc receptors for immunoglobulin G--a role in the pathogenesis of Guillain-Barré syndrome and multiple sclerosis. J Neuroimmunol. 2001;118(2):187–93.

20. León B, López-Bravo M, Ardavín C. Monocyte-derived dendritic cells formed at the infection site control the induction of protective T helper 1 responses against Leishmania. Immunity. 2007;26(4):519–31.

21. Zhou LJ, Tedder TF. CD14+ blood monocytes can differentiate into functionally mature CD83+ dendritic cells. Proc Natl Acad Sci U S A. 1996;93(6):2588–92.

22. Coyle PK. Detection and isolation of immune complexes in multiple sclerosis cerebrospinal fluid. J Neuroimmunol. 1987;15(1):97–107.

23. Coyle PK, Procyk-Dougherty Z. Multiple sclerosis immune complexes: an analysis of component antigens and antibodies. Ann Neurol. 1984;16(6):660–7.

24. Thompson AJ, Banwell BL, Barkhof F, Carroll WM, Coetzee T, Comi G, et al. Diagnosis of multiple sclerosis: 2017 revisions of the McDonald criteria. Lancet Neurol. 2018;17(2):162–73.

25. Tanaka M, Krutzik SR, Sieling PA, Lee DJ, Rea TH, Modlin RL. Activation of FcγRI on Monocytes Triggers Differentiation into Immature Dendritic Cells That Induce Autoreactive T Cell Responses. The Journal of Immunology. 2009;183(4):2349–55.

26. Kaskow BJ, Baecher-Allan C. Effector T Cells in Multiple Sclerosis. Cold Spring Harb Perspect Med. 2018;8(4):a029025.

27. van Langelaar J, Rijvers L, Smolders J, van Luijn MM. B and T Cells Driving Multiple Sclerosis: Identity, Mechanisms and Potential Triggers. Frontiers in Immunology. 2020;11(760).

28. León B, López-Bravo M, Ardavín C. Monocyte-Derived Dendritic Cells Formed at the Infection Site Control the Induction of Protective T Helper 1 Responses against <em>Leishmania</em>. Immunity. 2007;26(4):519–31.

29. Shortman K, Naik SH. Steady-state and inflammatory dendritic-cell development. Nat Rev Immunol. 2007;7(1):19–30.

30. Huang YM, Xiao BG, Ozenci V, Kouwenhoven M, Teleshova N, Fredrikson S, et al. Multiple sclerosis is associated with high levels of circulating dendritic cells secreting pro-inflammatory cytokines. J Neuroimmunol. 1999;99(1):82–90.

31. Pashenkov M, Teleshova N, Kouwenhoven M, Kostulas V, Huang YM, Soderstrom M, et al. Elevated expression of CCR5 by myeloid (CD11c+) blood dendritic cells in multiple sclerosis and acute optic neuritis. Clin Exp Immunol. 2002;127(3):519–26.

32. Laborde EA, Vanzulli S, Beigier-Bompadre M, Isturiz MA, Ruggiero RA, Fourcade MG, et al. Immune complexes inhibit differentiation, maturation, and function of human monocyte-derived dendritic cells. J Immunol. 2007;179(1):673–81.

33. Smed-Sörensen A, Moll M, Cheng T-Y, Loré K, Norlin A-C, Perbeck L, et al. IgG regulates the CD1 expression profile and lipid antigen-presenting function in human dendritic cells via FcγRIIa. Blood. 2008;111(10):5037–46.

34. Köller M, Zwölfer B, Steiner G, Smolen JS, Scheinecker C. Phenotypic and functional deficiencies of monocyte-derived dendritic cells in systemic lupus erythematosus (SLE) patients. International Immunology. 2004;16(11):1595–604.

35. Scheinecker C, Zwölfer B, Köller M, Männer G, Smolen JS. Alterations of dendritic cells in systemic lupus erythematosus: phenotypic and functional deficiencies. Arthritis Rheum. 2001;44(4):856–65.

36. Baba M, Ichinose K, Tamai M, Kawakami A, Ohyama K. Similarity of autoimmune diseases based on the profile of immune complex antigens. Rheumatol Int. 2019;39(2):323–5.

37. Ulvestad E, Williams K, Vedeler C, Antel J, Nyland H, Mørk S, et al. Reactive microglia in multiple sclerosis lesions have an increased expression of receptors for the Fc part of IgG. Journal of the Neurological Sciences. 1994;121(2):125–31.

